# Altered substance P signaling underlies perivascular sensory nerve dysfunction in inflammatory bowel disease

**DOI:** 10.1101/2020.04.29.066126

**Authors:** Charles E. Norton, Elizabeth A. Borgmann, Marcia L. Hart, Benjamin W. Jones, Craig Franklin, Erika M Boerman

## Abstract

**Objective:** Inflammatory Bowel Disease (IBD) is associated with cardiovascular disease risk and impaired intestinal blood flow, but the functional role of perivascular nerves that control vasomotor function of mesenteric arteries (MAs) perfusing the intestine is unknown in IBD. Because perivascular sensory nerves and their transmitters calcitonin gene-related peptide (CGRP) and substance P (SP) are important mediators of both vasodilation and inflammatory responses, our objective was to identify IBD-related deficits in perivascular sensory nerve function and vascular neurotransmitter signaling.

**Approach and Results:** In MAs from an IL-10^−/−^ mouse model, we found that IBD significantly impairs EFS-mediated sensory vasodilation and sensory inhibition of sympathetic vasoconstriction, despite decreased sympathetic nerve density and vasoconstriction. The MA content and EFS-mediated release of both CGRP and SP are slightly decreased with IBD, but IBD has different effects on each transmitter. CGRP nerve density, receptor expression, hyperpolarization and vasodilation are preserved with IBD. In contrast, SP nerve density and receptor expression are increased, and SP hyperpolarization and vasodilation are impaired with IBD. A critical finding is that blocking neurokinin 1 (SP) receptors restored EFS-mediated sensory vasodilation and enhanced CGRP-mediated vasodilation in MAs from IBD but not Control mice.

**Conclusions:** An aberrant role for the perivascular sensory neurotransmitter SP and its downstream signaling in MAs underlies vascular dysfunction with IBD. With IBD, SP signaling impedes CGRP-mediated sensory vasodilation, contributing to impaired blood flow. Substance P and NK1 receptors may represent an important target for treating vascular dysfunction in IBD.

## Introduction

Inflammatory Bowel Diseases (IBD) such as Crohn’s disease and ulcerative colitis are characterized by chronic aberrant immune responses and intestinal inflammation. IBD is comorbid with cardiovascular diseases including atherosclerosis^1^, arterial stiffening^2^, thromboembolism^3^ and heart failure^4^, which collectively point to an important role for vascular function in the pathogenesis and treatment if IBD. Although few studies have directly addressed changes in vascular function with IBD, disease manifestation is associated with impaired perfusion of the intestine^5, 6^ contributing to intestinal ischemia and pathology^7^.

Mesenteric arteries (MAs) which control blood flow to the gut, are widely utilized as a model of resistance arteries, but studies regularly ignore the role sensory innervation. MAs are enmeshed with both sympathetic and sensory nerves, which constrict and dilate MAs, respectively^8^. Perivascular sensory nerves further facilitate vasodilation via negative feedback inhibition of sympathetic neurotransmitter release^9, 10^ and participation in local axon reflexes^11^ in which inflammatory cytokines associated with IBD cause antidromic stimulation of sensory nerves and vasodilation^12^. Activated perivascular sensory nerves release co-packaged calcitonin gene-related peptide (CGRP) and substance P (SP), which both act as potent vasodilators^13, 14^. Released CGRP binds to a receptor complex consisting of the G-protein-coupled calcitonin receptor-like receptor, the essential chaperone receptor activity modifying protein (RAMP1) and the efficacy-enhancing receptor component protein. In SMCs, CGRP receptor activation promotes relaxation predominantly via cAMP and PKA-mediated activation of K^+^ channels^15, 16^ followed by decreased Ca^2+^ influx. The neurokinin SP acts as an endothelium dependent vasodilator^17^. Binding of SP to NK1 receptors on endothelial cells (ECs) mediates vasodilation via increased [Ca^2+^]_i_ and activation of endothelial nitric oxide synthase.

Impaired sensory nerve function has been implicated in decreased vasodilation and impaired vasomotor function in aging^18^, hypertension and diabetes^19^, underscoring the importance of sensory nerves in maintaining vascular homeostasis in both health and disease. Surprisingly, despite diminished vasodilator capacity^20^ and blood flow^5, 7, 21^ with IBD, the functional role of sensory nerves and their neuroeffector signaling pathways in IBD are unknown. Further, CGRP and SP have each been implicated as potential biomarkers for disease severity in human patients and have opposing effects in IBD^22^. Therefore, we tested the hypothesis that IBD causes perivascular sensory nerve dysfunction due at least in part to a shift in sensory nerve function from primarily CGRP-ergic to SP-ergic mechanisms.

## Materials and Methods

### Experimental animals

All experiments were performed in compliance with the Guide for the Care and Use of Laboratory Animals^23^ and were approved by the University of Missouri Animal Care and Use Committee. Male and female B6.129P2-IL10tm1Cgn/J (IL10^−/−^) mice, originally obtained from the Jackson Laboratory (Bar Harbor, ME) were bred and housed at the University of Missouri in a 12:12 light/dark cycle and received standard chow and water ad libitum. Prior to experimentation, mice were anesthetized via intraperitoneal injection of ketamine/xylazine (25mg/ml/2.5mg/ml). Following tissue dissection, mice were euthanized via cardiac exsanguination.

### Chemicals, Reagents, and Tools

Chemicals, antibodies and reagents used in this study are each listed in the Major Resources Table in the Supplemental Material along with their vendor, catalog information and concentration(s) used. Additional data are available upon request.

### Bacterial cultivation and inoculation

*H. hepaticus* (MU94) was grown on 5% sheep blood agar plates containing Brucella broth and 5% fetal bovine serum for 24-48h at 37°C in a microaerobic chamber with 90% N_2_, 5% H_2_, and 5% CO_2_. At 2 and 4 days post-weaning, the IBD group of mice received ∼10^8^ *H. hepaticus* (A_600_=1.0) suspended in 0.5 mL Brucella broth via gastric gavage. Non-gavaged littermates served as the Control group. Mice were then allowed to develop disease for 90 days before experimentation.

### Disease Scoring

To confirm the presence or absence of disease, cecum and colon tissue sections were collected and flushed with 10% neutral buffered formalin at euthanasia, formalin-fixed, paraffin-embedded, cut in 5-μm sections, and processed for hematoxylin and eosin staining. Fourteen random sections were selected from each group for disease scoring. To quantify disease, intestinal inflammation and hyperplasia were quantified post-hoc using an established blinded scoring system^24^.

### Pressure Myography

Mesentery and small intestine were placed in a chilled (4°C) dissection chamber containing Ca^2^+-free physiological salt solution (PSS) comprised of (in mM): 140 NaCl, 5 KCl, 1 MgCl_2_, 10 HEPES, 10 glucose (pH 7.4). A segment of intestine was pinned onto a block of transparent silicone (Dow Corning), and mesenteric arteries were hand-dissected away from surrounding mesentery and perivascular adipose.

Second-order arteries were transferred to the cannulation chamber (Warner Instruments) with a Wiretrol pipette, cannulated onto fire-polished ∼100μm glass pipettes, and secured with 11-0 suture. The pressure myography rig was mounted on an Olympus BX61W1 microscope. Arteries were pressurized to 70 mmHg and slowly heated to 37°C using an in-line heater (Warner Instruments). For all functional studies, arteries were superfused continuously at ∼5 ml/min with PSS (above) containing 2 mm CaCl_2_. To measure diameter, images were acquired using a 10X objective and a CCD camera (Basler) coupled to a computer to measure inner and outer diameters using edge-tracking software (LabView, National Instruments) customized by Dr. Michael J. Davis (University of Missouri).

Perivascular nerves were stimulated in an electrical field between the cannulation chamber’s two platinum electrodes connected to a stimulation isolation unit (SIU5, Grass) and square wave stimulator (S88, Grass). To measure sympathetic vasoconstriction and sensory vasodilation, electrical pulses (70 V, 2 ms) were delivered at 8 and 16 Hz to until a stable response diameter was achieved (<30 s). EFS-induced constrictions were measured in the presence and absence of the adrenergic agonist phenylephrine (1 μM), the sympathetic blocker guanethidine (10 μM), the sensory nerve blocker capsaicin (10μM), the CGRP receptor antagonist BIBN 4096 (1 μM) and the NK1 (SP) receptor antagonist CP 99994 (1 μM). To measure sensory inhibition of sympathetic vasoconstriction, arteries were stimulated at 1, 2, 4, 8 and 16 Hz in the absence ad presence of capsaicin (10 μM). The neurogenic nature of constrictions was verified by repeating the stimulations in the presence of the neurotoxin tetrodotoxin (TTX; 1 μM).

Concentration–response experiments were cumulative, with drug solutions added to the recirculating 20 ml superfusate. Drug concentrations to which vessels were exposed were: 0.1 nm - 1μm for CGRP and SP, 10 nm −10 μm for NE, 1 nm – 10 μm for ACh. Responses to CGRP, SP and ACh were measured following preconstriction with PE (1 μM). CGRP and SP responses were also measured in the presence of BIBN (1μM) and CP 99994 (1 μM). For all pressure myography and EFS studies, maximum diameter for each MA was obtained at the end of each experiment via superfusion with Ca^2+^-free PSS containing sodium nitroprusside (SNP, 10μm).

To measure structural and mechanical properties, cannulated arteries were equilibrated in Ca^2+^- free PSS containing 10 μM SNP at 37°C for 60 min. Intraluminal pressure (P) was lowered from 70 to 3 mmHg. Internal (D_i_) and external (D_e_) diameters were measured at 3 mmHg (D_0_) and after pressure steps in 20 mmHg increments from 20-120 mmHg. Values were used to calculate the following properties at each pressure as previously described in detail ^25, 26^.

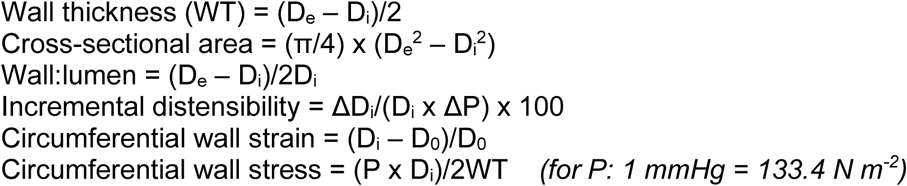

### Membrane potential recordings

Smooth muscle cell membrane potential (V_m_) was measured in pressurized arteries prepared as described above. To evaluate responses native to ECs, freshly isolated endothelial tubes were prepared via enzymatic dissociation as previously described and studied unpressurized at 32°C^27^. For both SMC and EC recordings, the electrophysiology platform was placed on an inverted microscope (Eclipse TS100, Nikon) and V_m_ was recorded using sharp microelectrodes backfilled with 2 M KCl (∼150-MΩ tip resistance) connected to an amplifier (AxoClamp 2B, Molecular Devices), data acquisition system (Digidata 1322A, Molecular Devices) and audible baseline monitor (model ABM-3, WPI) as previously described^27^. Baseline V_m_ was recorded for 10 min followed by increasing [CGRP] or [SP] (0.1nM – 1μM) added cumulatively in 5 min increments. Data were analyzed as 30 s averages during a stable V_m_ at each concentration.

### Immunofluorescence

Second-order MAs were pinned using 25 μm wire in a 24-well plate coated with Sylgard. For immunolabelling of perivascular nerves, MAs were pinned intact. For immunolabelling of RAMP1 and NK1, MAs were pinned *en face*. As previously described in detail^28^ arteries were fixed in 4% paraformaldehyde, blocked and permeabilized with PBS containing 1% BSA and 0.1% Triton X-100, and incubated overnight primary antibodies for tyrosine hydroxylase (TH), CGRP, SP, RAMP1 and/or NK1(see Supplemental Resources Table). Tissues were then blocked again, incubated in secondary antibodies, and mounted on slides using ProLong Gold with or without DAPI. Each experiment included no-primary controls that demonstrated a lack of non-specific fluorescence.

Slides were imaged using a Leica TCS SP8 confocal laser-scanning microscope (Leica Microsystems. Staining for TH-, CGRP-, and SP-containing nerves was visualized using a 25X water immersion objective and 1 μm Z-slices through one vessel wall. Staining for RAMP1 and NK1 was visualized using an HCX PL APO 60X glycerol immersion with a 2X optical zoom through the SMC and EC layers using 0.5 μm Z-slices. For all experiments, Control and IBD arteries were imaged using similar laser power and gain settings. Representative images were prepared by generating maximum z-projections through one vessel wall.

The density of PVNs was determined by quantifying the fluorescence area of CGRP, SP or TH staining on intact MA segments using ImageJ^29^. Briefly, maximum-intensity z-projections were converted to binary images with thresholds set to decrease background fluorescence. Fluorescence area fraction was then measured within the portion of the image occupied by the artery.

### Measurements of CGRP and SP

For arterial CGRP and SP content measurements, all first- and second-order MA segments from two mice were hand-dissected, pooled, weighed and frozen at −80°C. A tissue homogenate was made by mincing the arteries with a scalpel blade and homogenizing with a hand-held homogenizer in 400 µl of 1X PBS. Tissue homogenates underwent centrifugation of 5 min at 12,000 x g at 4°C to pellet debris. ELISA measurements from each MA sample were normalized to the wet weight of segments. CGRP and SP release were measured from the same mice as arterial content. A long (>1mm) first-order mesenteric artery was cannulated, pressurized and heated as previously described. The vessel length, inner diameter and outer diameter were then measured and recorded. Superfusion was then stopped, 500 μl PSS was added to the cannulation chamber, and the artery was subjected to EFS (60 V, 2 ms, 1 min). The 500 μl bath solution was removed and set aside. A second artery was then cannulated as before and stimulated in the same 500 μl bath solution collected from artery #1. This solution was then collected and frozen at −80°C. ELISA values for each sample were then normalized to the total adventitial volume of artery stimulated for that sample, calculated as follows:

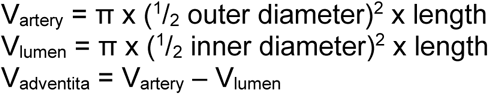

CGRP and Substance P levels for all samples were measured using a CALCA ELISA (mouse, Aviva Systems Biology #OKCD00726) and Substance P ELISA (mouse, Biomatik #EKC37868) kit. These kits were selected for the content and release measurements due to their higher sensitivity range compared to the kits used for serum measurements.

For serum CGRP and SP measurements, whole blood was collected via cardiac puncture from both control and IBD animals and allowed to clot at room temperature for 30 min. Blood was then centrifuged for 15 min at 2,000 x g at 4°C and serum supernatant collected and stored at −80°C. Undiluted serum samples were analyzed for CGRP and SP using Fluorescent ELISA kits (Phoenix Pharmaceuticals) n=19 (Control) and 20 (IBD) mice.

### Statistical Analysis

All data are reported as means ± SEM. Statistical significance was determined at P< 0.05. n values represent the number of biological replicates used in each experiment. Statistical analysis was completed using Graphpad Prism 7. All concentration-response, electrical field stimulation and vessel wall mechanics data were analyzed by 2-way repeated measures ANOVA followed by Bonferroni’s *post hoc* tests. Disease scoring, nerve density and ELISA data were analyzed by Student’s unpaired t tests.

### Chemicals, Reagents, and tools

Chemicals and reagents used in this study are each listed in the Major Resources Table along with their vendor, catalog information and concentration(s) used. The table also includes vendors and catalog numbers for unique tools and equipment.

## Results

### IL10^−/−^ + H. hepaticus model of IBD

IBD mice (IL10^−/−^ mice gavaged with *H. hepaticus*) developed significant epithelial hyperplasia and severe inflammation throughout the cecum and colon associated with higher histological disease scores compared to Control mice (IL10^−/−^ littermates not treated with *H. hepaticus*) (Figure 1). Thus, IL-10 deficiency alone in these mice does not cause IBD.

**Figure 1.**
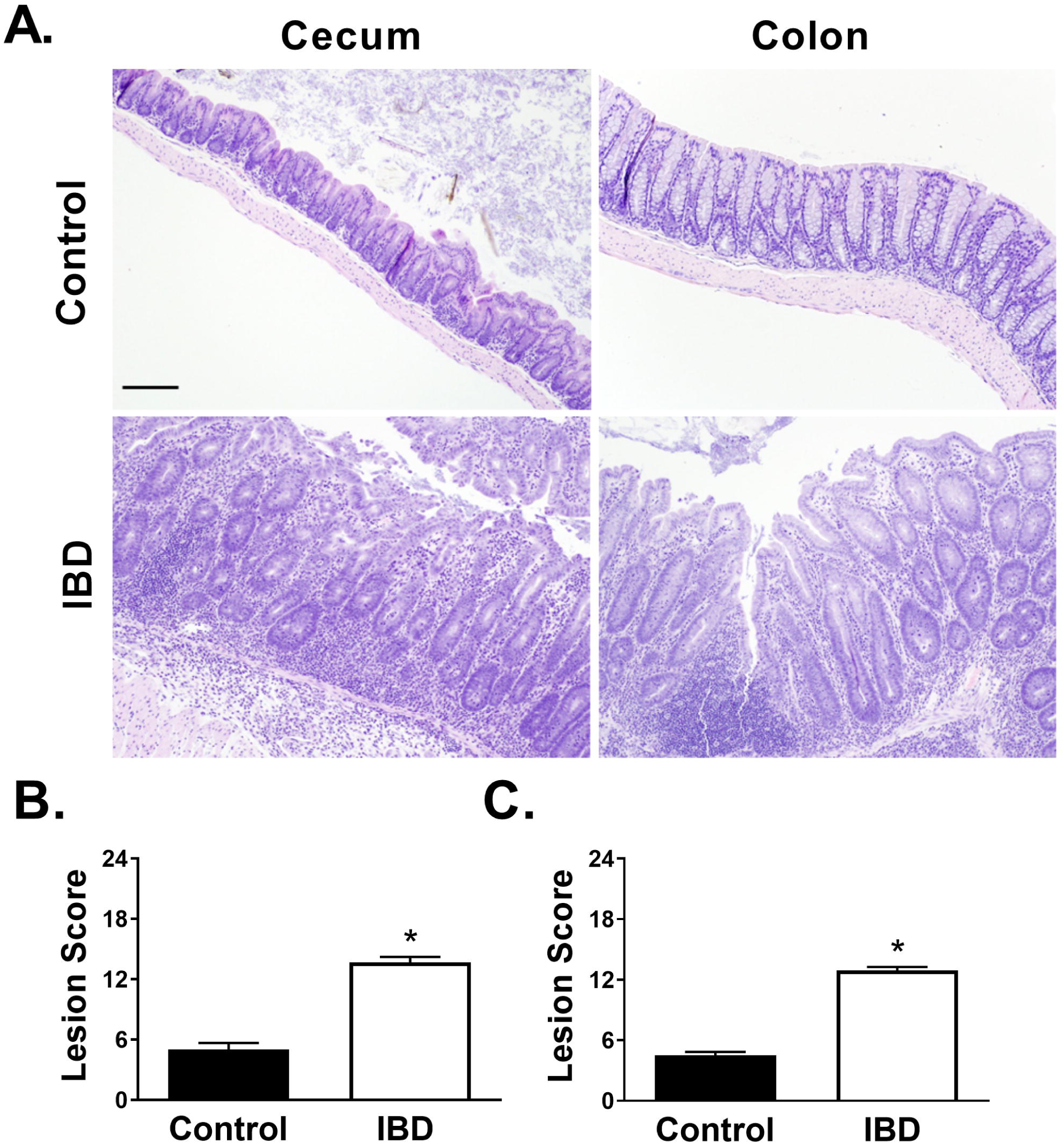
*Helicobacter hepaticus*-induced colitis in IL10^−/−^ mice. **A)** Representative images of H&E stained cecum and colon sections from Control (IL10^−/−^) or IBD (*H. hepaticus*-inoculated IL10^−/−^) mice. **B and C)** Tissue lesion scores (Mean ± SE, range 0-24) in **B)** cecum and **C)** colon tissues. n=14 per group; * = P<0.05 vs Control. Scale bar = 50 μm

### IBD impairs sympathetic vasoconstriction, sensory vasodilation and sympathetic inhibition

To measure perivascular sympathetic nerve function, vasomotor responses in cannulated MAs were measured following EFS. Electrically-induced constrictions in MAs were decreased in IBD vs Control arteries at both 8 and 16 Hz (Figure 2A) indicating deficits in perivascular sympathetic nerve function. To assess the role of sensory vasodilation, sympathetic nerves were blocked with guanethidine (10 μM), and arteries were preconstricted to a similar degree with PE (Control 44 ± 5% and IBD 41 ± 2%). Vasodilation was significantly impaired in IBD vs Control arteries at both 8 and 16Hz (Figure 2B). Sensory mediation of each dilation was confirmed via its elimination by pre-treatment with the sensory nerve blocker capsaicin (CAP; 10 μM, Figure 2C). Sensory inhibition of sympathetic vasoconstriction increased constriction in Control arteries (Figure 2D) but was absent in IBD arteries (Figure 2E), suggesting a lack of sympathetic inhibition by sensory nerves IBD. The neurotoxin TTX (1 μM) blocked all EFS-induced constrictions in both groups at 1-16 Hz (Figure 2G), confirming that EFS responses were nerve-mediated.

**Figure 2.**
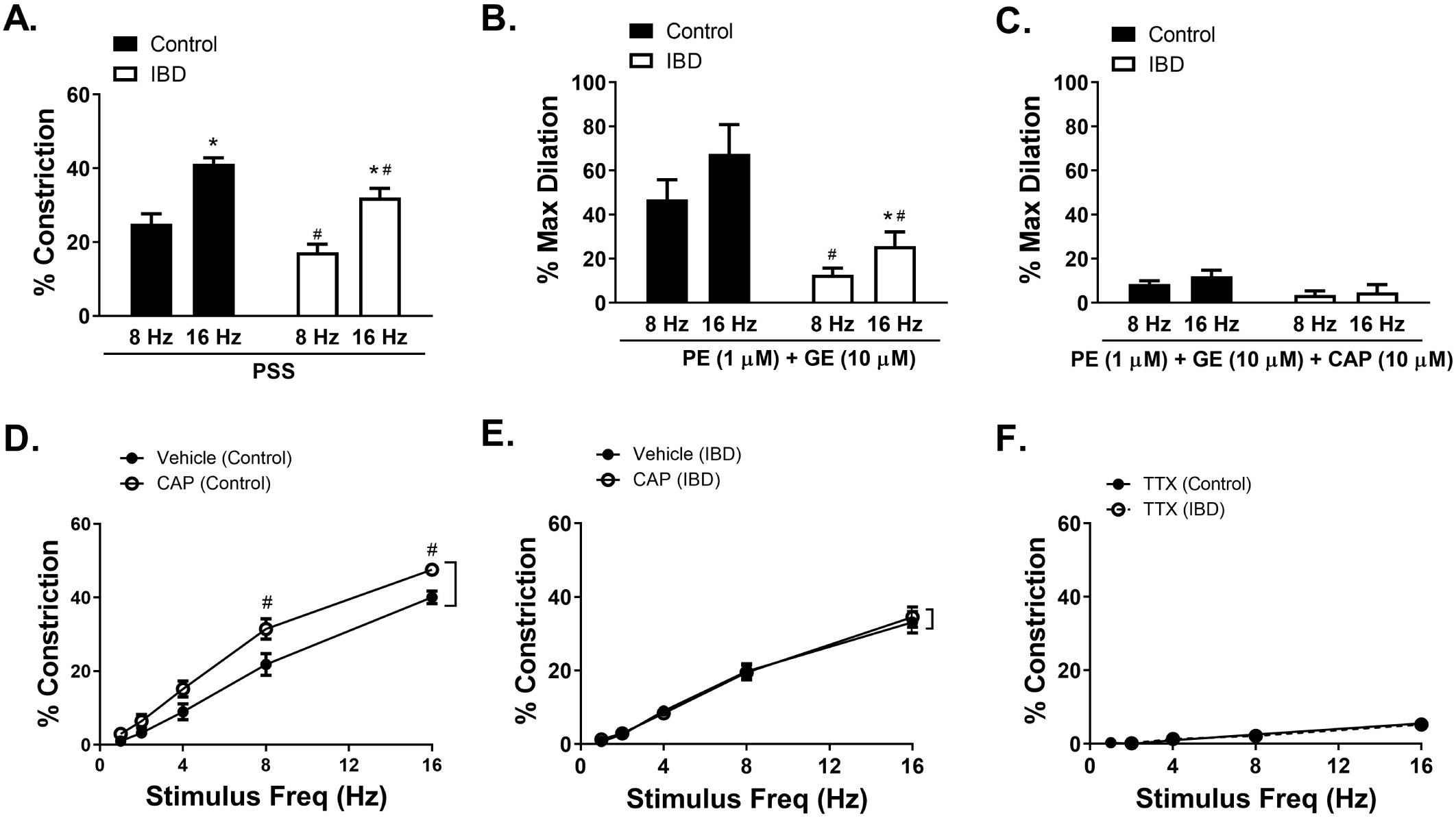
IBD impairs sympathetic constriction, sensory dilation, and sensory inhibition of sympathetic nerves. **A)** Electrical Field Stimulation (EFS) induced (8 and 16 Hz) sympathetic constrictions of isolated, cannulated and pressurized arteries. **B)** EFS-induced (8 and 16 Hz) sensory dilation of phenylephrine (PE, 1 μM) and guanethidine (GE, 10 μM) treated arteries **C)** that is blocked by capsaicin (CAP; 10 μM). **D and E)** Sensory inhibition of sympathetic constriction in **E)** Control and **F)** IBD arteries, shown as difference in EFS-induced (1-16Hz) constriction in the presence and absence of CAP (10 μM). **F)** Tetrodotoxin (TTX) inhibition of EFS-induced (1-16 Hz) constriction. Data are mean ± SE % constriction or % max dilation. * = P<0.05 16 vs 8 Hz; # = P<0.05 IBD vs Control. n=5-8 per group.

### Vasomotor, structural and mechanical properties of mesenteric arteries

As a general index of smooth muscle and endothelial function, cumulative concentration-response curves were generated for the adrenergic vasoconstrictor norepinephrine (NE, 1 nM-10 μM) and the EC-dependent vasodilator Acetylcholine (ACh, 1 nM-10 μM. NE produced similar LogEC_50_ (Control: −6.62 ± 0.05; IBD: −6.90 ± 2.40) and constrictions in both groups (Supplemental Figure IA), whereas IBD desensitized PE-preconstricted arteries to ACh, with LogEC_50_ values increasing from −7.42 ± 0.09 to −6.7 7± 0.08 despite similar maximum dilations (Supplemental Figure IB). Thus, deficits in the overall ability of MAs to constrict and dilate likely does not underlie the significant impairments in perivascular nerve function. Passive pressure-diameter curves were created for cannulated arteries to determine whether structural and mechanical changes contribute impaired sympathetic vasoconstriction and sensory vasodilation with IBD. IBD was not associated with significant changes in inner diameter, outer diameter, wall thickness, cross-sectional area, wall-lumen ratio, incremental distensibility, circumferential wall stress, or circumferential wall strain at pressure intervals from 20-120 mmHg (Supplemental Figure IIA-H).

### Perivascular nerve density and neurotransmitter content/release

Mesenteric arteries were immunolabeled for CGRP, SP, and TH to determine whether altered or imbalanced nerve density underlies impaired sensory nerve function with IBD. CGRP nerve density was unchanged while SP nerve density was modestly increased in IBD arteries. In contrast, TH nerve density significantly decreased in IBD arteries (Figure 3), suggesting that sensory nerve dysfunction in IBD persists despite a relative decrease in the presence of sympathetic nerves. To determine if the content or release of sensory neurotransmitters changed in MAs with IBD, ELISA measurements were made on samples of MA homogenate and post-EFS bath solution. There was a nonsignificant decrease (P≈0.10) in CGRP and SP content in mesenteric artery homogenates from IBD vs Control arteries (Figure 4A-B). Consistent with effects on content, CGRP and SP released into bath solution following EFS of cannulated arteries had a tendency to be lower in IBD vs Control samples (Figure 4C-D). however, circulating serum CGRP were similar between groups (Figure 4E-F).

**Figure 3.**
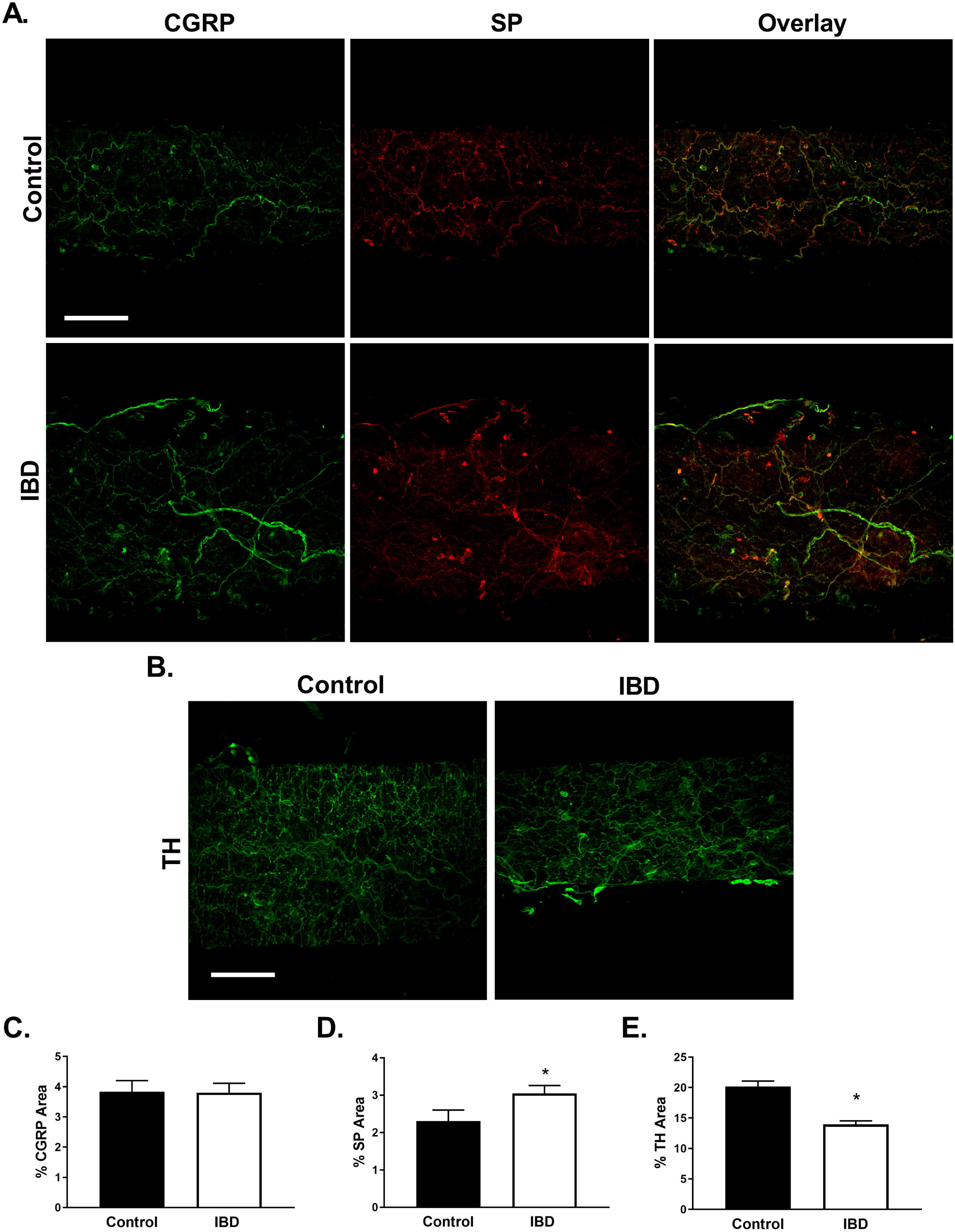
Perivascular nerve density is altered with IBD. **A and B)** Representative maximum z-projections through the adventitia showing labeling for **A)** Calcitonin Gene-related Peptide (CGRP, left), Substance P (SP, center), and CGRP-SP overlay (right) in Control (Upper) and IBD (Lower) arteries **B)** Tyrosine Hydroxylase (TH) in Control (left) and IBD (right) arteries. Scale bar = 100 μm. **C-E)** Quantitation of nerve density for **C)** CGRP **D)** SP and **E)** TH. Data are mean ± SEM for percent fluorescence within the vessel wall. * = P<0.05 vs Control. n=11-21 vessel segments from 4 mice per group.

**Figure 4.**
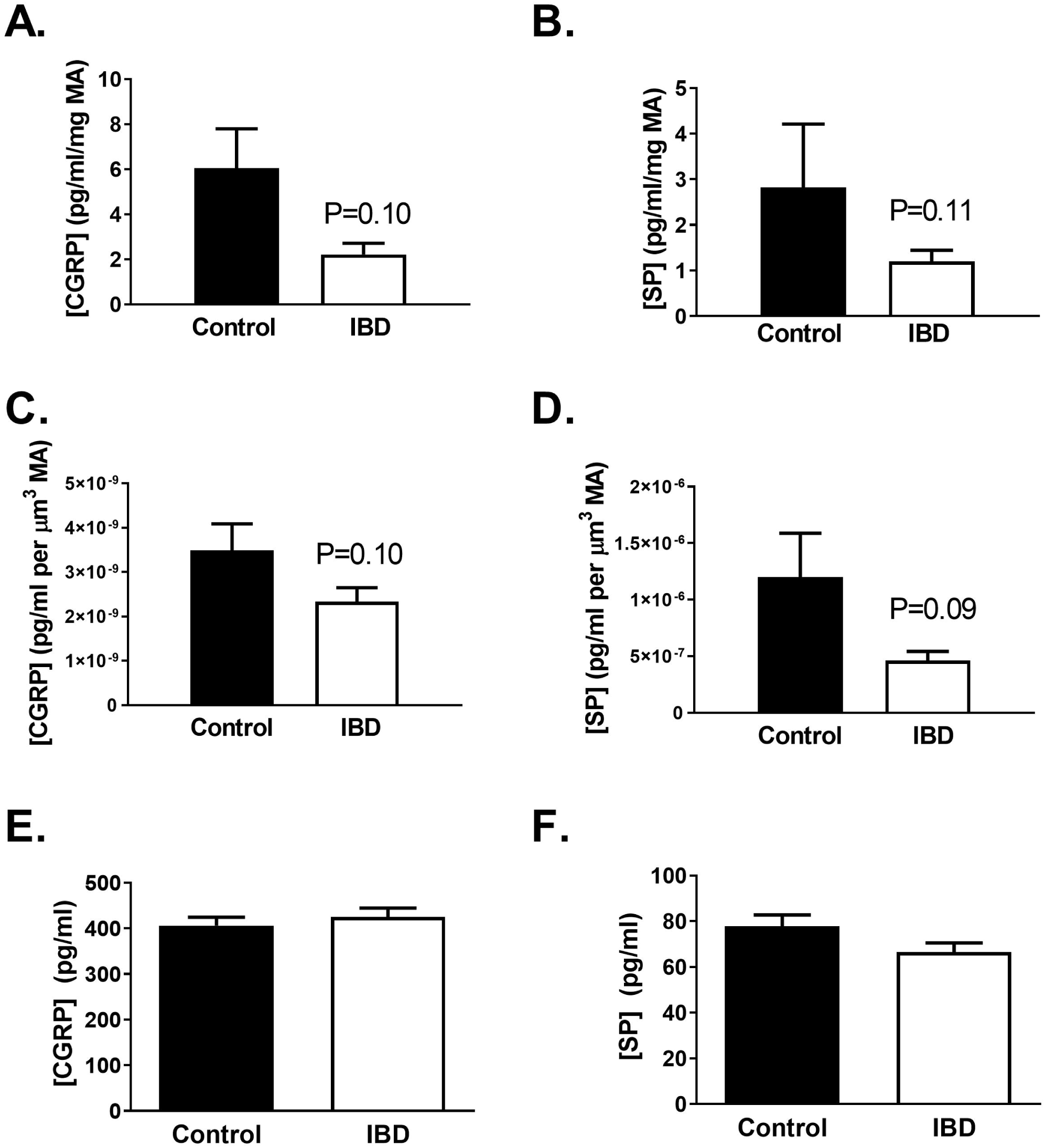
IBD decreases sensory neurotransmitter content and release from MAs. IBD was associated with trends toward (**A and B**) decreased CGRP and SP content in intact MAs and (**C and D**) decreased EFS-mediated CGRP and SP release into tissue bath measured by ELISA. n=3 replicates from 6 mice. **E and F)** Serum SP and CGRP were similar in Control vs IBD mice. N=19-20.

### CGRP and SP receptor function and expression

Because EFS-mediated sensory vasodilation was impaired with IBD, we conducted concentration-response experiments for the sensory neuropeptides CGRP and SP (0.1 nM-1 μM) as an index of their downstream receptor function. CGRP-mediated dilations were similar in Control and IBD arteries, with equal potency (LogEC_50_ Control −8.45 ± 0.16 and IBD −8.13 ± 0.27 respectively) and a trend toward decreased efficacy with IBD (Max dilations: 73 ± 2% to 57 ± 4) (Figure 5A). SP dilation was significantly impaired with IBD; despite similar potency (LogEC_50_: −8.33 ± 0.15 and −8.11 ± 0.28), maximum dilations were significantly reduced (Figure 5B). The observed decreases in SP but not CGRP-mediated dilation correspond to changes in V_m_. CGRP caused concentration-dependent hyperpolarization of SMCs (Figure 5C) and ECs (Figure 5E) that was similar between Control and IBD. Hyperpolarization of both SMCs (Figure 5D) and ECs (Figure 5F) to SP was significantly attenuated with IBD. Expression of RAMP1 (CGRP receptor protein) was unchanged in both SMCs and ECs with IBD. NK1 receptors were not expressed SMCs of either group but had increased immunolabeling in ECs with IBD (Figure 6), potentially upregulated in response to decreased nerve content and release. Taken together, these data suggest that changes in SP receptor function contribute to impaired perivascular sensory nerve function with IBD.

**Figure 5.**
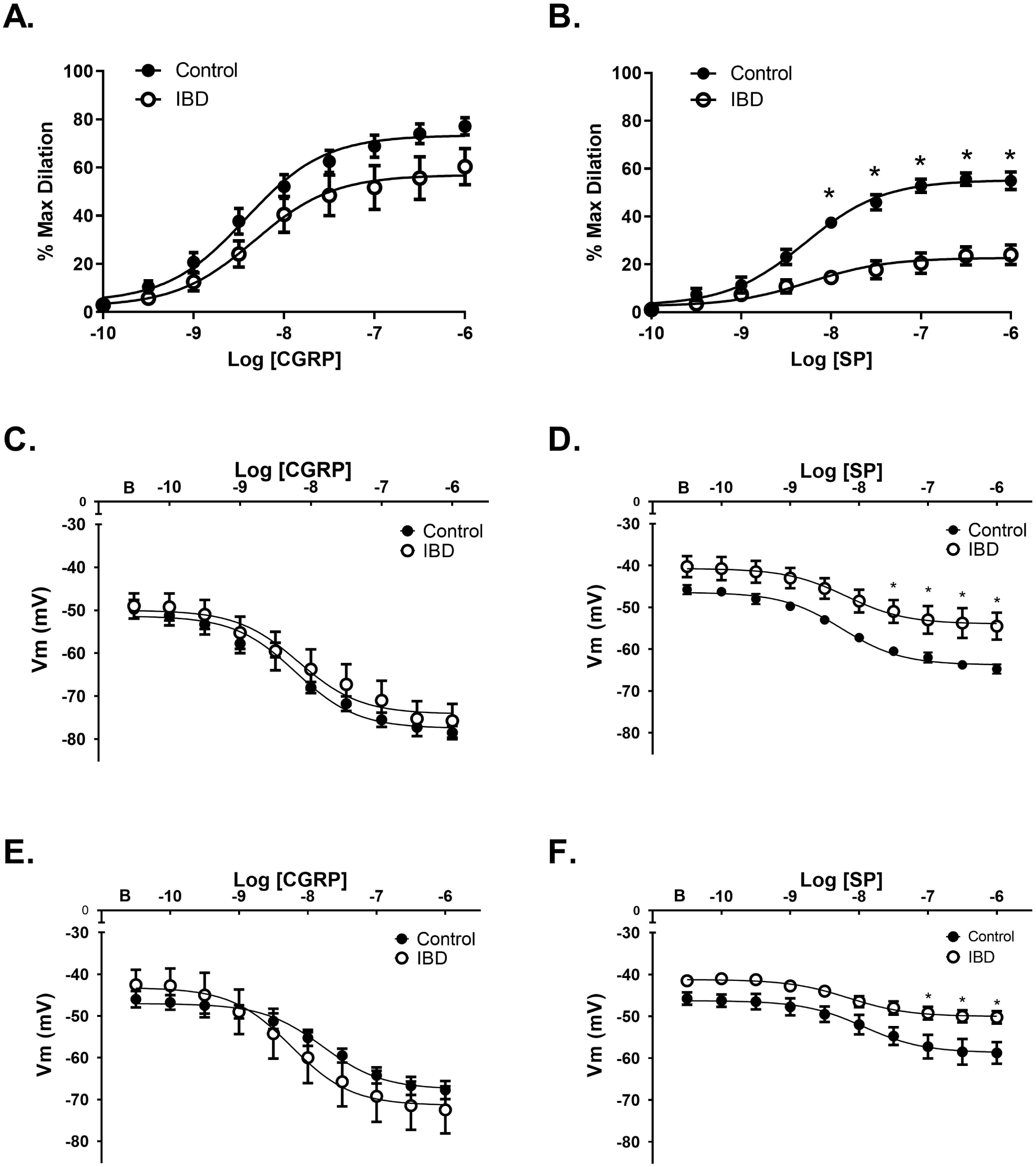
IBD impairs dilation and hyperpolarization to exogenous SP but not CGRP. **A and B)** Vasodilation and **C-D)** membrane hyperpolarization of isolated, cannulated and pressurized arteries exposed to cumulative concentrations of **A and C)** CGRP and **B and D)** SP. **E and F)** Membrane hyperpolarization of intact EC tubes exposed to cumulative concentrations of **E)** CGRP and **F)** SP. Data are mean ± SE for % max dilation (n=9-11) and V_m_(n=4-5) per group. *=P<0.05 vs Control. B=baseline V_m_.

**Figure 6.**
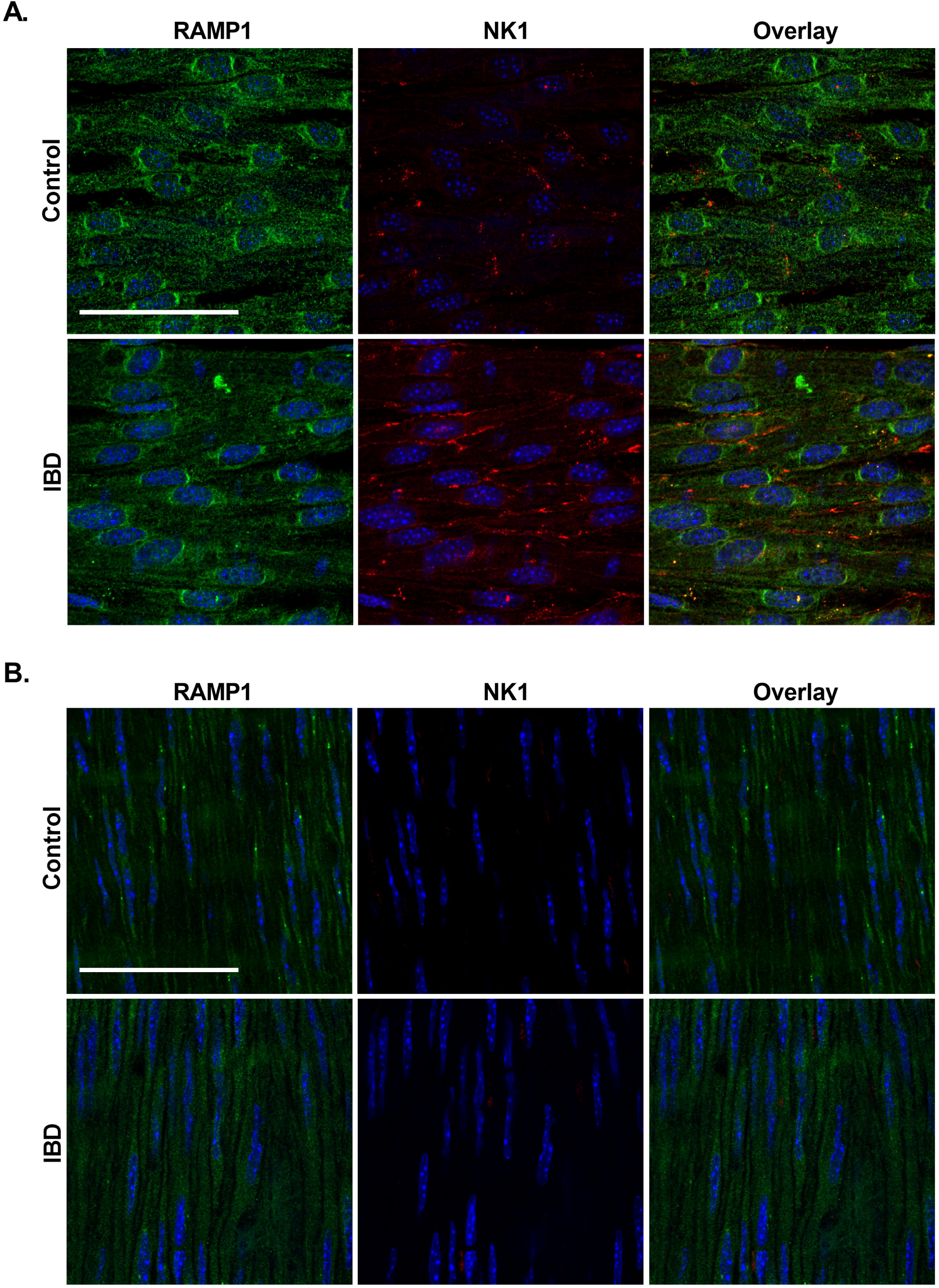
IBD alters NK1 but not RAMP1 expression and localization in mesenteric arteries. Panels are representative maximum z-projections through the **A)** endothelial and **B)** smooth muscle cell layers of *en face* mesenteric arteries showing labeling for CGRP receptors (Receptor Activity Modifying Protein (RAMP1, left) Neurokinin 1 receptors (NK1, center), and RAMP1-NK1 overlay (right) in Control (Upper) and IBD (Lower) arteries. All images include a DAPI nuclear stain. Scale bar = 50μm. n=16 vessel segments from 4 mice per group.

### IBD alters role of nerve-released SP but not CGRP in mesenteric arteries

To determine whether the relative contribution of CGRP and SP to sensory vasodilation changes with IBD, we measured sensory dilations of preconstricted (PE) sympathetically-blocked (GE,) MAs to EFS in the absence and presence of CGRP and/or SP receptor blockade with BIBN 4096 and CP 99994 respectively. As in previous EFS experiments (Figure 2), total sensory vasodilation was significantly impaired in IBD vs Control at 8Hz and 16Hz. Blockade of the CGRP-ergic contribution with BIBN 4096 decreased sensory dilation of Control arteries ∼60% at both 8 and 16Hz. Subsequent blockade of the SP-ergic contribution with CP99994 (BIBN+CP) did not significantly decrease sensory dilation vs BIBN alone at 8Hz or 16Hz (Figure 7A). Reversing the order of application, dilation of Control arteries was not significantly decreased by CP at 8Hz, but concurrent CGRP receptor blockade (CP+BIBN) decreased sensory dilation at both 8Hz and 16Hz (Figure 7B). These data suggest that in healthy MAs, CGRP is predominantly responsible for sensory dilation, with a lesser role for SP.

**Figure 7.**
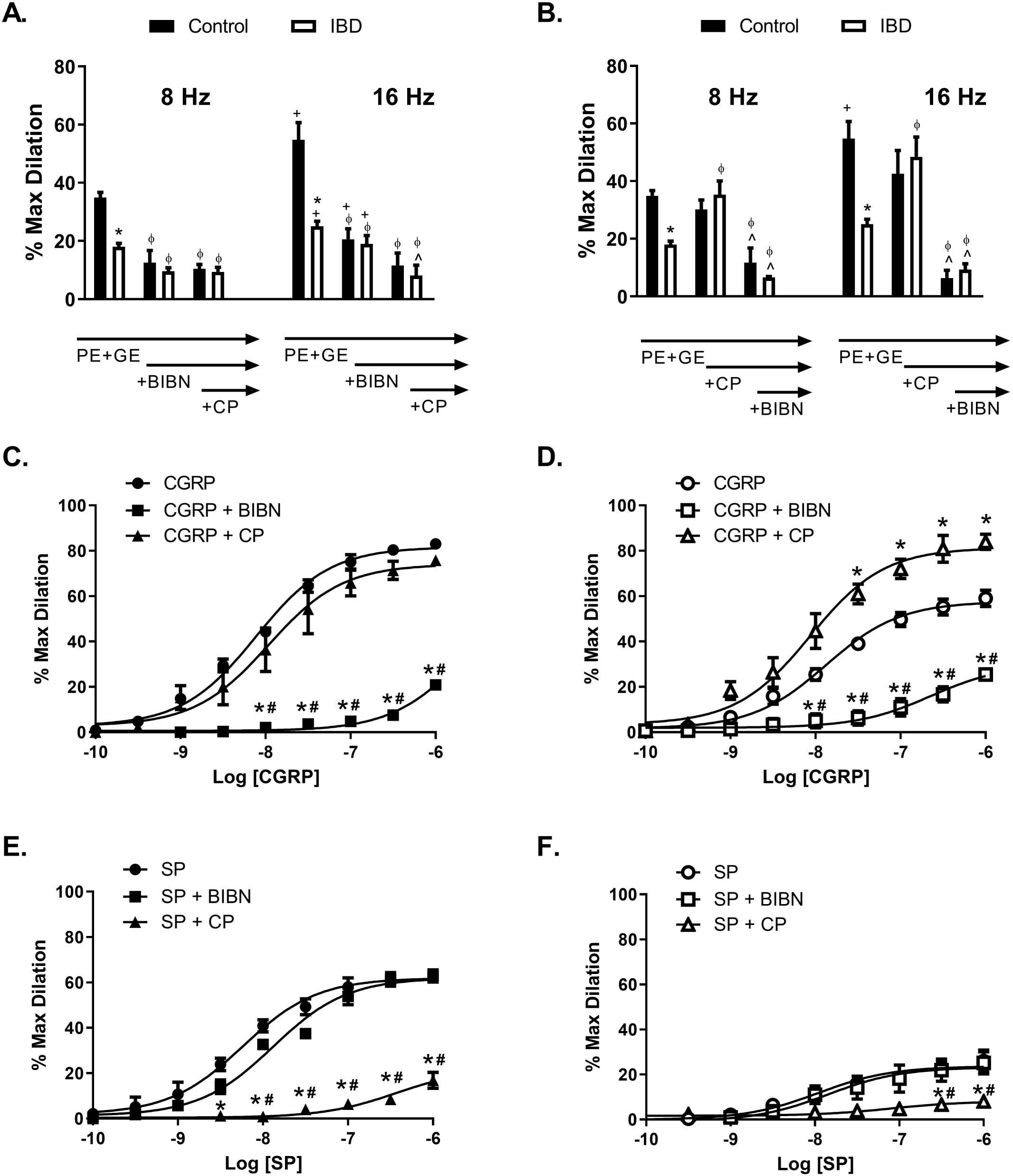
Respective roles of CGRP and SP receptor activation in EFS- and neuropeptide-induced vasodilation. **A and B)** EFS-induced (8 and 16 Hz) sensory dilation of PE (1μM) and GE (10 μM)-treated arteries in the presence of **A)** the CGRP receptor antagonist BIBN 4096 (1 μM) and BIBN + the NK1 antagonist CP99994 (1 μM) or **B)** CP + BIBN. Data are mean ± SE % maximum dilations. *=P<0.05 Control vs IBD; ϕ=P<0.05 vs PE+GE; +=P<0.05 16 vs 8 Hz; ^=P<0.05 vs PE+GE+BIBN (A) or PE+GE+CP (B). N=4-6 per group. **C-F)** Vasodilation of isolated, cannulated and pressurized **C and E**) Control and **D and F)** IBD arteries exposed to cumulative concentrations of **C and D)** CGRP and **E and F)** SP in the absence and presence of BIBN or CP. *=P<0.05 vs CGRP or SP alone; #=P<0.05 vs CGRP+CP or SP+BIBN. n=3-6 per group.

In IBD arteries, BIBN blocked ∼45% of sensory dilation at 8Hz and ∼25% at 16 Hz. Subsequent addition of CP (BIBN+CP) did not change sensory dilation vs BIBN alone at 8Hz but significantly attenuated dilation at 16 Hz (Figure 7A), consistent with an increased role for SP in sensory dilation with IBD. Reversing antagonist order had a distinct and surprising effect in IBD arteries: NK1 blockade with CP *restored* the previously-impaired sensory vasodilations at both 8Hz and 16Hz (Figure 7B). Subsequent addition of BIBN (CP+BIBN) significantly reduced dilations at 8Hz and 16Hz (Figure 7B). Thus, during IBD, SP appears to impede CGRP-mediated dilation, leading to significant decreases in EFS-induced sensory vasodilation.

To verify both the specificity of the antagonists and the altered role of SP during IBD, we generated concentration-response curves to CGRP and SP in the absence and presence of each antagonist. CGRP-mediated vasodilation was effectively eliminated in the presence of BIBN in both Control and IBD (Figure 7C-D) arteries, and SP-mediated vasodilation was eliminated in the presence of CP in both groups (Figure 7E-F). CP did not affect CGRP dilation of Control arteries (Figure 7C) but, consistent with EFS experiments, increased CGRP-mediated dilation of IBD arteries (max dilation 59±4% to 83±3%) (Figure 7D). BIBN had no effect on SP dilation in Control (or IBD (Figure 7E-F) arteries. These data demonstrate SP-mediated attenuation of CGRP vasodilation in MAs with IBD.

## Discussion

Using an immune-driven IL-10^−/−^ mouse model, we found that IBD impairs perivascular sympathetic and sensory nerves on MAs, with the largest deficits in EFS-mediated sensory vasodilation and sensory inhibition of sympathetic vasoconstriction. SP-but not CGRP-containing nerve density is increased in MAs with IBD, and the density of opposing TH-containing sympathetic nerves decreases. IBD decreases storage and release of both CGRP and SP despite unchanging circulating transmitter. Substance P and its downstream signaling appears to play a predominant role in mediating vascular dysfunction with IBD. Arteries from IBD mice fail to normally hyperpolarize and dilate to exogenously-applied SP, and expression of SP, but not CGRP receptor proteins is increased in MAs with IBD. Further, blocking NK1 receptors restores both EFS-mediated sensory dilation and CGRP-mediated vasodilation in IBD but not control MAs. These data suggest that with IBD, SP impairs CGRP signaling, thereby decreasing sensory vasodilation and blood flow to the intestine.

Perivascular nerves represent a novel target for the treatment of IBD. They control blood flow to the intestines, which is impaired in IBD^5, 6^. Perivascular nerves also regulate intestinal function, as nerves on MAs enter the intestinal wall and terminate in the submucosal and myenteric plexuses that influence motility and secretion^30^. Unfortunately, the role of sympathetic and sensory innervation of the intestine and associated vasculature in clinical and experimental IBD remains controversial. Nerve density measurements and denervation studies have been commonly used to elucidate the role of sympathetic and sensory nerves during IBD. Global sympathectomy can ameliorate and exacerbate colitis, depending on the model used (see summary in^31^). Presently, we found a decrease in sympathetic nerve density (Figure 3), which is consistent with modestly decreased TH axons in gut submucosal arteries in 2,4,6-Trinitrobenzenesulfonic acid (TNBS) colitis^32^ but in contrast to unchanged TH nerves in human submucosal arteries^33^ and increased TH nerves in human MAs^34^ with IBD. Sensory denervation exacerbates colitis in female oxazolone-induced^35^, TRPV1^−/−^ Dextran Sulfate Sodium (DSS)- induced^36^ mouse models and some TNBS-induced rat models^37, 38^ but improves colitis in a different TNBS rat model^39^ and DSS and TNBS mouse models^40^. We report increased SP and unchanged CGRP nerve density on MAs with IBD (Figure 3), similar to reports in human submucosal arteries^34^. In light of decreased sympathetic nerve density and vasoconstriction with IBD (Figure 2), the increase in SP nerves would be expected to improve sensory dilation and sympathetic inhibition; that they do not either supports an aberrant role for adventitial SP nerves or upregulation to combat decreased downstream signaling.

This is the first study to directly examine perivascular sensory nerve function in the mesenteric vasculature with IBD. EFS-induced sensory vasodilation of MAs was significantly impaired with IBD, as was sensory inhibition of sympathetic vasoconstrictions (Figure 2). Our data suggest that this is not due to increased sympathetic opposition (Figures 2-3) but rather a change in SP and NK1 function during IBD. Dysfunction of NK1 receptors with IBD is highlighted by diminished SP-mediated dilation and hyperpolarization of MAs (Figure 5) despite increased expression of endothelial NK1 receptors (Figure 6). Substance P signaling contributes to the pathogenesis of multiple diseases^41-43^, including IBD, where SP knockout mice are protected against colitis^44^. Crohn’s patients have significantly increased expression of NK1 receptors in vasculature, enteric neurons and lymphoid aggregates of the large intestine^45^. Colon NK1 expression also increases in experimental colitis, where NK1 blockade decreases colonic inflammation^46-48^. Thus, SP and its receptors are important contributors to both mucosal inflammation and vascular dysfunction during IBD. Compared to CGRP, Substance P is largely pro-inflammatory: serum SP increases in IBD patients, with circulating levels paralleling disease severity and decreasing during remission^49^. This contrasts with our finding of unchanged serum SP and CGRP (Figure 4E-F) and suggests the NK1 upregulation and aberrant SP-mediated signaling observed in this study do not arise from increased whole-body SP. In contrast to SP, CGRP is anti-inflammatory in many tissues^50, 51^ and participates in immune responses by decreasing the release of proinflammatory cytokines from multiple cell types^52, 53^. In patients with IBD, CGRP is decreased in the intestinal mucosa, and the magnitude of this decrease is correlated with the severity of IBD symptoms^54^. CGRP knockout mice are susceptible to colitis^44^, further implicating an important protective role for CGRP in IBD. In this study vascular CGRP nerves and receptors persist during IBD (Figures 3 and 6) and continue to dilate MAs in response to exogenous CGRP (Figure 5) despite significantly impaired EFS-mediated sensory vasodilation (Figure 2). This suggests that SP signaling in ECs is the critical factor underlying sensory nerve dysfunction in IBD.

More significant than impaired SP-mediated dilation is its apparent ability to alter CGRP signaling with IBD. We found that EFS-induced sensory dilations were restored to control levels in the presence of NK1 blockade (Figure 7A-B). NK1 blockade also improved CGRP-mediated vasodilation with IBD (Figure 7C-D), suggesting that perivascular SP becomes functionally pro-contractile with IBD by interfering with CGRP-mediated vasodilation. The underlying mechanism of this SP effect is unclear in our study but is consistent with research showing that co-injection of SP with CGRP converts prolonged CGRP-mediated increases in blood flow to transient effects in human skin^55^. Those effects were linked to SP-induced mast cell release of a CGRP-degrading protease. Mast cells accumulate in the human gastrointestinal tract during IBD, and their release of mast cell protease exacerbates colitis in mouse DSS and TNBS models^56^. Whether mast cells or other immune cells infiltrate the vascular adventitia during IBD and how they may affect perivascular nerve function is unknown. Nevertheless, SP and its associated signaling pathways appear to play a central role in perivascular nerve dysfunction with IBD. Future work will focus understanding the multifaceted roles of SP in the vasculature in health and IBD in order to target its detrimental effects (interference with CGRP vasodilation, exacerbated inflammation) while preserving its ability to facilitate vasodilation and intestinal perfusion.

## Acknowledgments

None

## Abbreviations

ACH: Acetylcholine
CAP: Capsaicin
CGRP: Calcitonin Gene-related Peptide
DSS: Dextran Sulfate Sodium
EC: Endothelial Cell
EFS: Electrical Field Stimulation
GE: Guanethidine
IL-10: Interleukin 10
MA: Mesenteric Artery
NK1: Neurokinin 1
PE: Phenylephrine
RAMP1: Receptor Activity Modifying Protein
SMC: Smooth Muscle Cell
SP: Substance P
TH: Tyrosine Hydroxylase
TNBS: 2,4,6-Trinitrobenzenesulfonic acid solution,
TTX: Tetrodotoxin

## Highlights

- Inflammatory Bowel Disease causes significant impairment of perivascular sensory nerve mediated vasodilation and inhibition of sympathetic vasoconstriction in mesenteric arteries that perfuse the intestine.
- Despite impaired substance P vasodilation, SP containing nerve density and NK1 receptor expression increase in MAs with IBD; CGRP dilation, nerve density and receptor expression are not changed.
- Blocking NK1 receptors restores sensory vasodilation in MAs and increases CGRP-mediated vasodilation, indicating that SP interference with CGRP signaling underlies impaired sensory vasodilation with IBD.

