## Supplemental Materials for "Altered substance P signaling underlies perivascular sensory nerve dysfunction in inflammatory bowel disease"

**SUPPLEMETNAL MATERIAL**

**Running title**: Perivascular nerve dysfunction in IBD

**Corresponding author**:

Erika M. Boerman, PhD

Department of Medical Pharmacology and Physiology

MA415 Medical Science Building - 1 Hospital Drive

University of Missouri

Columbia, MO 65212

**Keywords**: mesenteric arteries, CGRP, substance P, vasodilation

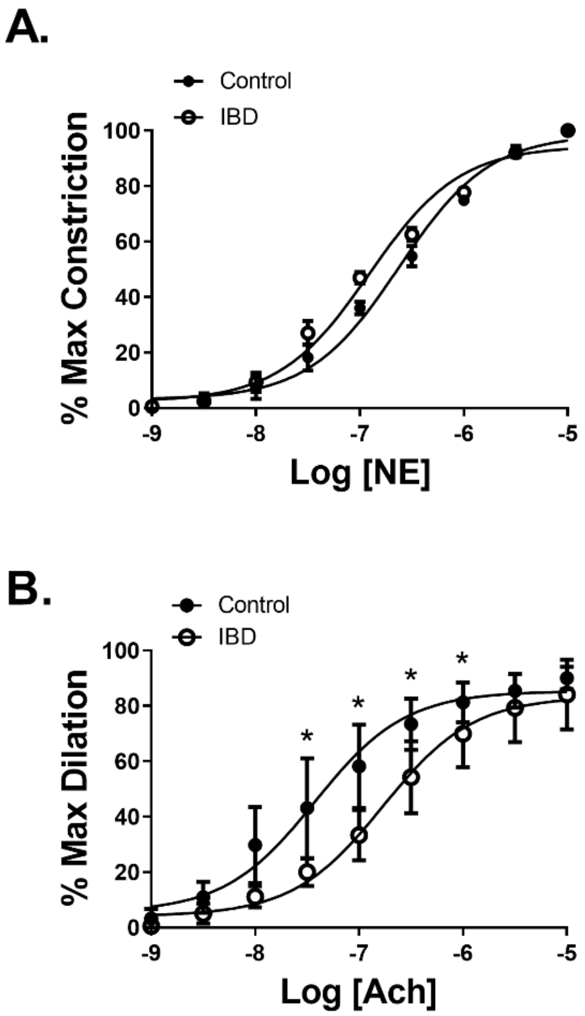

**Supplementary Figure I. IBD desensitizes arteries to ACh but not NE**. **A)** Data are mean ± SEM for percent maximum dilation (obtained in 0 $Ca^{2+}$ PSS + 10 μM SNP) of isolated, cannulated pressurized arteries preconstricted with phenylephrine (1 μM) and treated with cumulative concentrations of acetylcholine (Ach, 1 nM-10 μM). **B)** Data are mean ± SEM for percent maximum constriction (normalized to 100%) of isolated, cannulated pressurized arteries treated with cumulative concentrations of norepinephrine (NE, 1 nM-10 μM). *=P<0.05 vs Control. n=6 per group

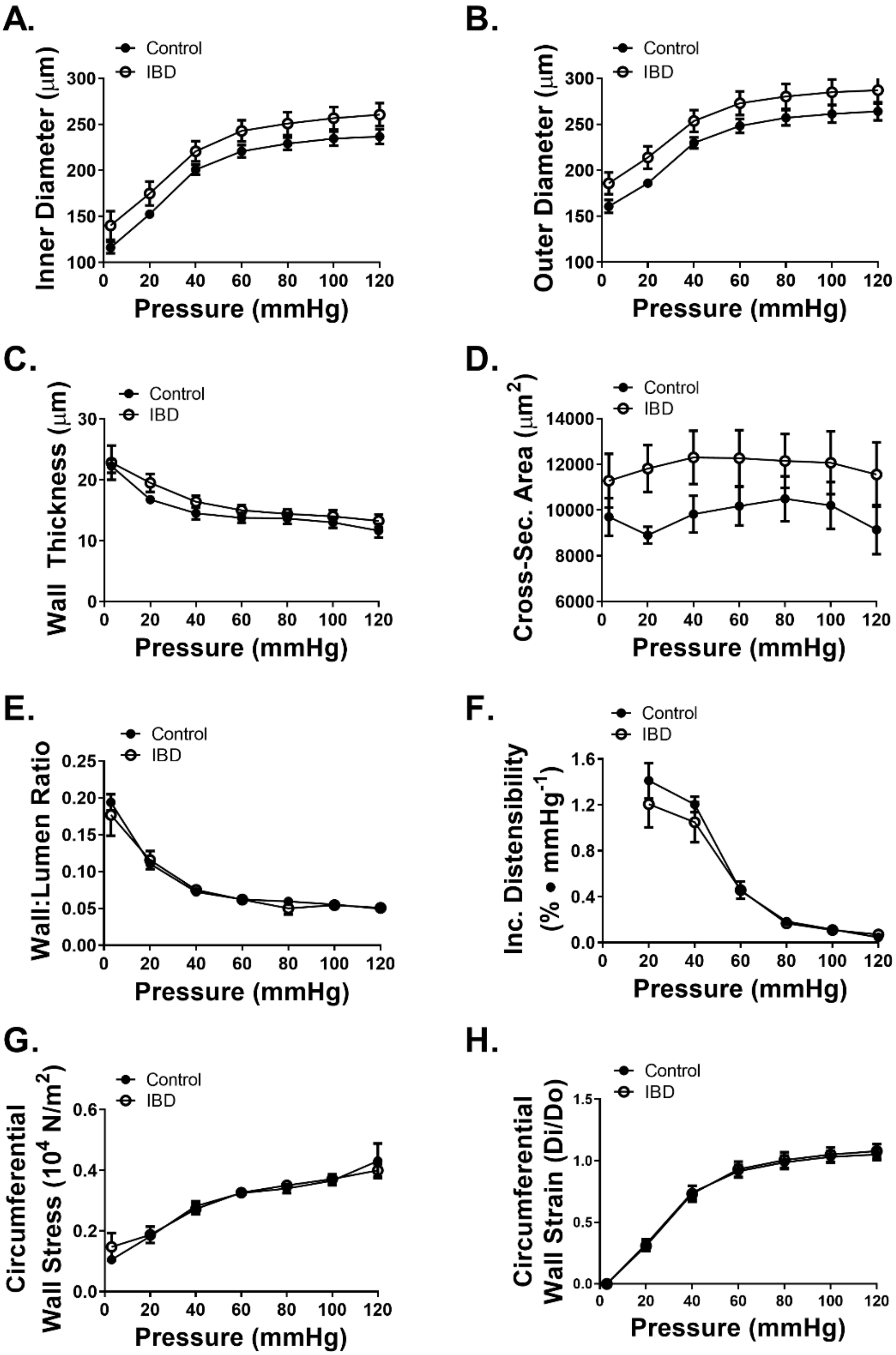

A.
Control
IBD
Inner Diameter (μm)
Pressure (mmHg)
B.
Control
IBD
Outer Diameter (μm)
Pressure (mmHg)
C.
Control
IBD
Wall Thickness (μm)
Pressure (mmHg)
D.
Control
IBD
Cross-Sec. Area (μm2)
Pressure (mmHg)
E.
Control
IBD
Wall:Lumen Ratio
Pressure (mmHg)
F.
Control
IBD
Inc. Distensibility (% • mmHg-1)
Pressure (mmHg)
G.
Control
IBD
Circumferential Wall Stress (104 N/m2)
Pressure (mmHg)
H.
Control
IBD
Circumferential Wall Strain (Di/Do)
Pressure (mmHg)

**Supplementary Figure II. IBD does not alter structural or mechanical properties of mesenteric arteries. A)** Inner diameter (μm), **B)** Outer diameter (μm), **C)** Wall thickness, **D)** Cross-sectional area, **E)** Wall:lumen, **F)** Incremental distensibility, **G)** Circumferential wall stress, and **H)** Circumferential wall strain are similar in Control and IBD arteries. Data are mean ± SEM for each variable. n=6 per group.

**Major Resources Tables**

### Animals (*in vivo* studies)

| Species | Vendor or Source | Background Strain | Sex |
|---|---|---|---|
| Murine | The Jackson Laboratory | B6.129P2-*Il10*$^{tm1Cgn}$/J, stock no: 002251 | Male & female |

### Animal breeding

| | Species | Vendor or Source | Background Strain | Other Information |
|---|---|---|---|---|
| **Parent - Male** | Murine | The Jackson Laboratory | B6.129P2-*Il10*$^{tm1Cgn}$/J | Stock no: 002251 |
| **Parent - Female** | Murine | The Jackson Laboratory | B6.129P2-*Il10*$^{tm1Cgn}$/J | Stock no: 002251 |

### Antibodies

| Target antigen | Vendor or Source | Catalog # | Working concentration | Lot # |
|---|---|---|---|---|
| Calcitonin gene-related peptide | MilliporeSigma | PC205L | 1:250 | 3093330 |
| Substance P | Abcam | ab14184 | 1:250 | GR3214181-3 |
| Tyrosine Hydroxylase | MilliporeSigma | T2928 | 1:250 | 067M4877V |
| Receptor activity-modifying protein 1 | Santa Cruz | sc-11379 | 1:250 | *now discontinued |
| Neurokinin receptor 1 (Tacr1) | Invitrogen | PA1-32229 | 1:250 | |
| Goat-anti-rabbit Alexa Fluor 488 (used w/ CGRP, RAMP1) | ThermoFisher Scientific | A-11034 | 1:500 | 1937195 |
| Goat-anti-mouse Alexa Fluor 633 (used w/ SP, TH) | ThermoFisher Scientific | A-21050 | 1:500 | |
| Chicken-anti-rat Alexa Fluor 647 (used w/ NK1) | ThermoFisher Scientific | A-21472 | 1:500 | 1263041 |

### ELISA kits

| Target antigen | Vendor or Source | Catalog # | Standard Curve $R^2$ | Lot # |
|---|---|---|---|---|
| CGRP | Aviva Systems Biology | OKCD00726 | 0.999 | KC0595 |

| | | | |
|---|---|---|---|
| SP | Biomatik | EKC37868 | 1.000 |
| CGRP | Phoenix Pharmaceuticals | FEK-015-09 | 0.989 |
| SP | Phoenix Pharmaceuticals | FEK-061-05 | 0.999 |

**Drugs and Reagents**

| Name | Vendor or Source | Catalog Number | Concentration used |
|---|---|---|---|
| Capsaicin | MilliporeSigma | M2028 | 10 µM |
| Tetrodotoxin | Tocris | 1078 | 1 µM |
| (R)-(-)-Phenylephrine hydrochloride | MilliporeSigma | P6126 | 1 µM |
| Guanethidine sulfate | BOC Sciences | 645-43-2 | 10 µM |
| Calcitoni~~o~~n gene-related peptide | Anaspec | AS-20682 | 0.1 nm – 1 µm |
| Substance P | Anaspec | AS-24280 | 0.1 nm – 1 µm |
| Acetylcholine chloride | MilliporeSigma | A6625 | 1 nm – 10 µm |
| BIBN 4096 | Tocris | 4561 | 1 µM |
| CP 99994 | Tocris | 3417 | 1 µM |
| Sodium nitroferricyanide (III) | MilliporeSigma | 228710 | 10 µM |
| Collagenase from *Clostridium histolyticum* | MilliporeSigma | C8051 | |
| Elastase | MilliporeSigma | 324682 | |

**Relevant Supplies and equipment**

| Name | Vendor or Source | Catalog Number | Other Information |
|---|---|---|---|
| ProLong Gold mounting media with DAPI | ThermoFisher Scientific | P36941 | |
| BBL Brucella Broth | BD Difco | 211088 | |
| Fetal Bovine Serum | MilliporeSigma | F0926 | |
| 5% Sheep blood agar plates | Hardy Diagnostics | A10 | |
| 11-0 monofilament nylon | Ashaway Line & Twine Manufacturing Company | 5300001 | 7 ply |
| Pipette glass | Warner Instruments | 64-0790 & 64-0781 | Standard wall with filament model G120F-4 & Thin wall model G120T-4 |

| | | | |
|---|---|---|---|
| Wiretrol II pipette | Drummond Scientific Company | 5-000-2100 | For transferring vessel segments |
| Pressure myography/EFS chamber | Warner Instruments | 640240R2 | Model RC-27NE2 |
| Sylgard 184 | MilliporeSigma | 761036 | |
| CCD camera | Basler | scA1600-28fm | |
| In-line heater | Warner Instruments | 640104 | Model SHM-6 |
| Temperature controller | Warner Instruments | 642401 | Model TC-344C |
| Stimulus Isolation Unit | Grass instruments | SIU5 | Out of production |
| Square wave generator | Grass Instruments | S88 | Out of production |
| Scissors for *en face* prep | Fine Science Tools | 15000-03 | 2 mm cutting edge |
| Stainless steel 27 μm pinning wire | Ebay | https://ebay.to/2LvK0qu | |
